# A diet-independent zebrafish model for NAFLD recapitulates patient lipid profiles and offers a system for small molecule screening

**DOI:** 10.1101/2022.04.11.487858

**Authors:** Manoj K Singh, Rohit Yadav, Akash Kumar Bhaskar, Shantanu Sengupta, Chetana Sachidanandan

## Abstract

Non-alcoholic Fatty Liver Disease (NAFLD) or pathological hepatic lipid overload, is considered to affect obese individuals. However, NAFLD in lean individuals is prevalent, especially in South Asian population. The pathophysiology of lean NAFLD is not well understood and most animal models of NAFLD use the high-fat diet paradigm. To bridge this gap, we have developed a diet-independent model of NAFLD in zebrafish. We have previously showed that chronic systemic inflammation causes metabolic changes in the liver leading to hepatic fat accumulation in an IL6 overexpressing (IL6-OE) zebrafish model. In the present study, we compared the hepatic lipid composition of adult IL6-OE zebrafish to the controls and found an accumulation of saturated triacylglycerols and a reduction in the unsaturated triacylglycerol species reminiscent of NAFLD patients. Zebrafish is an ideal system for chemical genetic screens. We tested whether the hepatic lipid accumulation in the IL6-OE is responsive to chemical treatment. We found that PPAR-gamma agonist Rosiglitazone, known to reduce lipid overload in the high fat diet models of NAFLD, could ameliorate the fatty liver phenotype of the IL6-OE fish. Rosiglitazone treatment reduced the accumulation of saturated lipids and showed a concomitant increase in unsaturated TAG species in our inflammation-induced NAFLD model. Our observations suggest that the IL6-OE model can be effective for small molecule screening to identify compounds that can reverse hepatic lipid accumulation, especially relevant to lean NAFLD.

## Introduction

Non-alcoholic fatty liver disease (NAFLD), or the pathological overaccumulation of fat in the liver is one of the most insidious chronic liver diseases globally. It affects 20-30% of the global population and the prevalence of NAFLD is on an upward trend over the last many years(1). Currently, there are no effective therapies or treatments for NAFLD other than lifestyle and diet related changes(2,3). Obesity and high body adiposity are considered as triggers for fatty liver(4,5). Thus, most rodent models of NAFLD rely on high fat or high fructose diet to induce fatty liver(6). However, there is a rising incidence of obesity-independent NAFLD, especially in South Asians for which the causes are unclear (7). To date, no single animal model is reported to recapitulate the complete spectrum of human NAFLD, although features similar to human NAFLD have been observed in some models that are helpful in understanding disease pathophysiology. Inflammation is prominently observed during NAFLD (8); however, the mechanism behind inflammation-induced steatosis is still poorly understood.

Recently, we described a diet-independent zebrafish fatty liver model where chronic inflammation induced by IL6 overexpression (IL6-OE) causes metabolic changes in the liver leading to hepatic fat accumulation (9). In all NAFLD models studied, excessive accumulation of neutral lipid, especially triglycerides in the liver, is associated with dysregulation of glucose and lipid metabolism that leads to inflammation and further deterioration of the liver leading to fibrosis and cirrhosis(10). In the inflammation-induced fatty liver model in zebrafish we found dysregulated expression of glycolytic and lipid metabolism genes, but the pattern was distinct from the high fat/fructose diet models(9).

In this study, we have performed lipidomics studies on the adult IL6-OE and control livers. Our studies revealed that the IL6-OE fish liver accumulates more saturated and mono-unsaturated triacylglycerols (TAGs) and shows a reduction in the polyunsaturated triacylglycerol species reminiscent of NAFLD patients (11). Zebrafish is an ideal vertebrate model for chemical screening(12,13). We took advantage of this facet of the zebrafish model and performed a small targeted chemical screen for compounds that can ameliorate the lipid accumulation in the IL6-OE fish larvae. We found that Rosiglitazone, a PPAR-gamma agonist, could ameliorate the hepatic fat accumulation in IL6-OE larvae. This likely reverses the downregulation of PPAR signaling components observed in the IL6-OE. Adult IL6-OE zebrafish liver treated with rosiglitazone showed reduction in the accumulation of saturated lipids and a concomitant increase in unsaturated TAGs. This is in concordance to previously reported pre-clinical studies, where rosiglitazone treatment reduced hepatic fat accumulation in high fat diet models of NAFLD and type 2 diabetes (14,15).

## Results

### IL6 overexpression leads to excess hepatic fat accumulation in zebrafish liver

We used a double transgenic *Tg (myl7:GAL4-VP16)::(UAS:HsaIL6)* zebrafish model (henceforth known as IL6-OE) to study the fatty liver condition. In this model, *myl7* promoter drives the expression of *GAL4-VP16* in cardiac tissue and this induces the overexpression of secreted human IL6 from the heart. In our previous studies with this zebrafish line, we demonstrated that systemic overexpression of IL6 causes excess fat accumulation in the adult liver(9). We discovered that a dysregulation of the glucose metabolism is associated with the fatty liver phenotype. The IL6-OE zebrafish would thus be an ideal system to screen small molecules that can reverse the fatty liver phenotype. To characterize the embryonic and larval stages of the IL6-OE transgenic line so that they can be used for the chemical genetic screening, we determined the expression of IL6 in the early stages by RNA in-situ hybridization. Expression of human IL6 mRNA was evident as early as 24hpf in the IL6-OE embryos but not in the control (*Tg (myl7:GAL4-VP16))* (Fig 1A,B). Quantitative RT-PCR of 5dpf larvae revealed that the transgenic animals had very high expression of human IL6 (Ct values of 28.3) compared to controls (Ct value of 41.7) (Fig. 1C). Systemic induction of IL6 signaling was evidenced by the 1.6-fold induction of *socs3,* a STAT3 target gene (Fig. 1D). Systemic IL6 is known to induce hepcidin in the liver(16) and we found this to be true in the IL6-OE embryos as well with a 3.6-fold induction of hepcidin (Fig. 1E).

**Figure 1.**
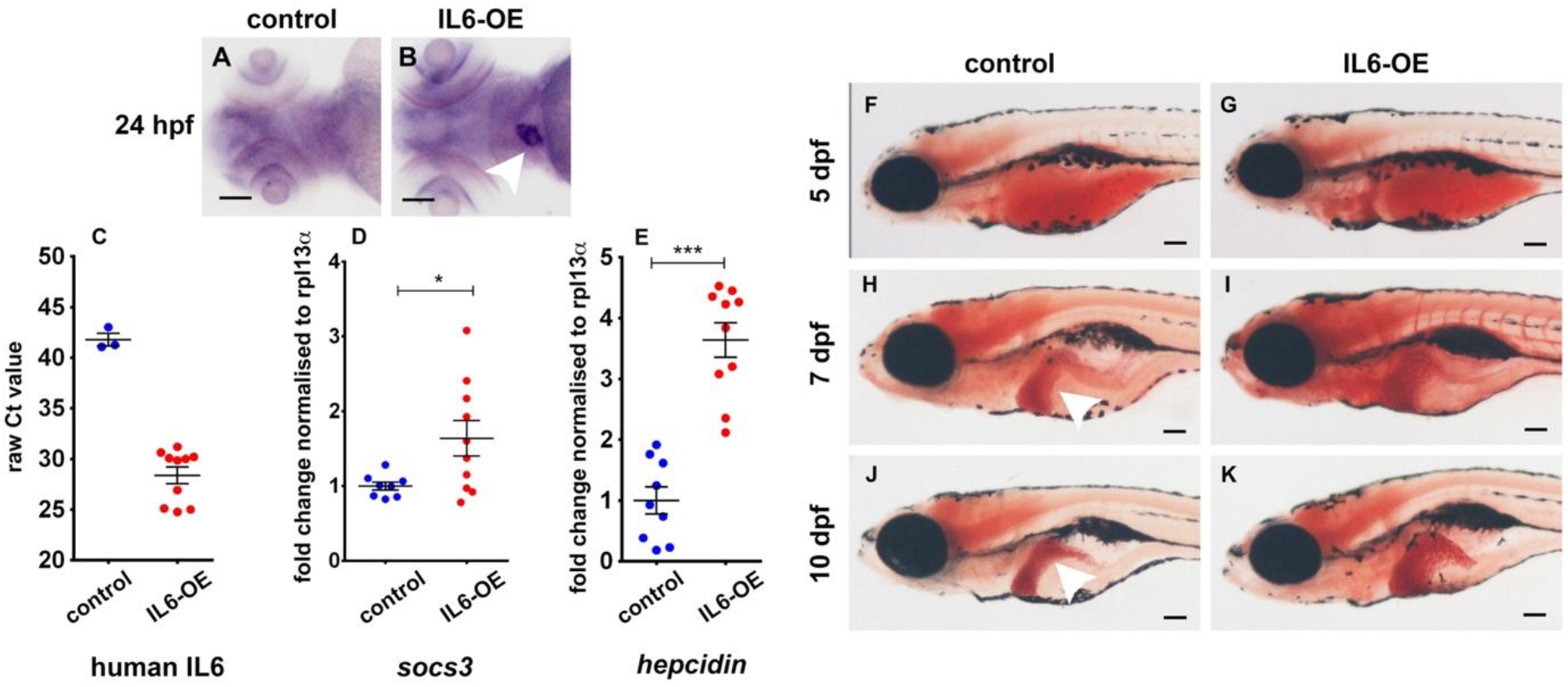
IL6 overexpression leads to fat accumulation in zebrafish liver. (A, B) RNA in situ hybridization shows expression of human IL6 mRNA in the heart of 1dpf IL6-OE zebrafish larvae. C) qRT-PCR of human IL6 shows higher expression in IL6-OE compared to control. y-axis shows raw Ct value. (D, E) qRT-PCR of *socs3* and *hepcidin* shows significant induction in IL6-OE compared to control. (F-K) Oil red O staining of 5dpf, 7dpf and 10dpf zebrafish larvae shows increase in fat accumulation in the IL6-OE liver compared to controls. Scale bar is 10μm.

Since, our previous study has shown that IL6-OE fish accumulates fat in the liver(9), we used Oil red O (ORO) stain to detect fat accumulation in liver in the zebrafish larvae and found that at 5dpf there was no significant difference between control and IL6-OE. However, by 7dpf there was a clear increase in hepatic fat as well as liver size in the IL6-OE larvae compared to the control. This difference was much more pronounced in the 10dpf larvae (Fig. 1F-K).

### Suppression of lipogenesis, cholesterol metabolism, β-oxidation and glycolytic enzymes in IL6-OE zebrafish larvae

Our previous studies on IL6-OE adult zebrafish liver had shown distinct expression pattern of signature genes in comparison to previously established high fat diet (HFD) models of hepatic steatosis(9). To check the expression pattern of genes involved in lipid and glucose metabolism, we performed qRT-PCR on 7dpf control and IL6-OE larvae We observed a downward trend in expression of lipogenesis genes in larvae with significant down regulation of *srebp1, pparab, pparg* and *cd36* (Fig. 2A). Cholesterol metabolism gene *hmgcs1* was also significantly downregulated while *hmgcra* showed a downward trend (Fig. 2B). β-oxidation leads to breakdown of fatty acid in the mitochondria and peroxisomes; we found significant downregulation of the peroxisomal β-oxidation gene *acox3* (Fig. 2C). These patterns of gene expression were very similar to those observed in the IL6-OE adult liver (9).

**Figure 2.**
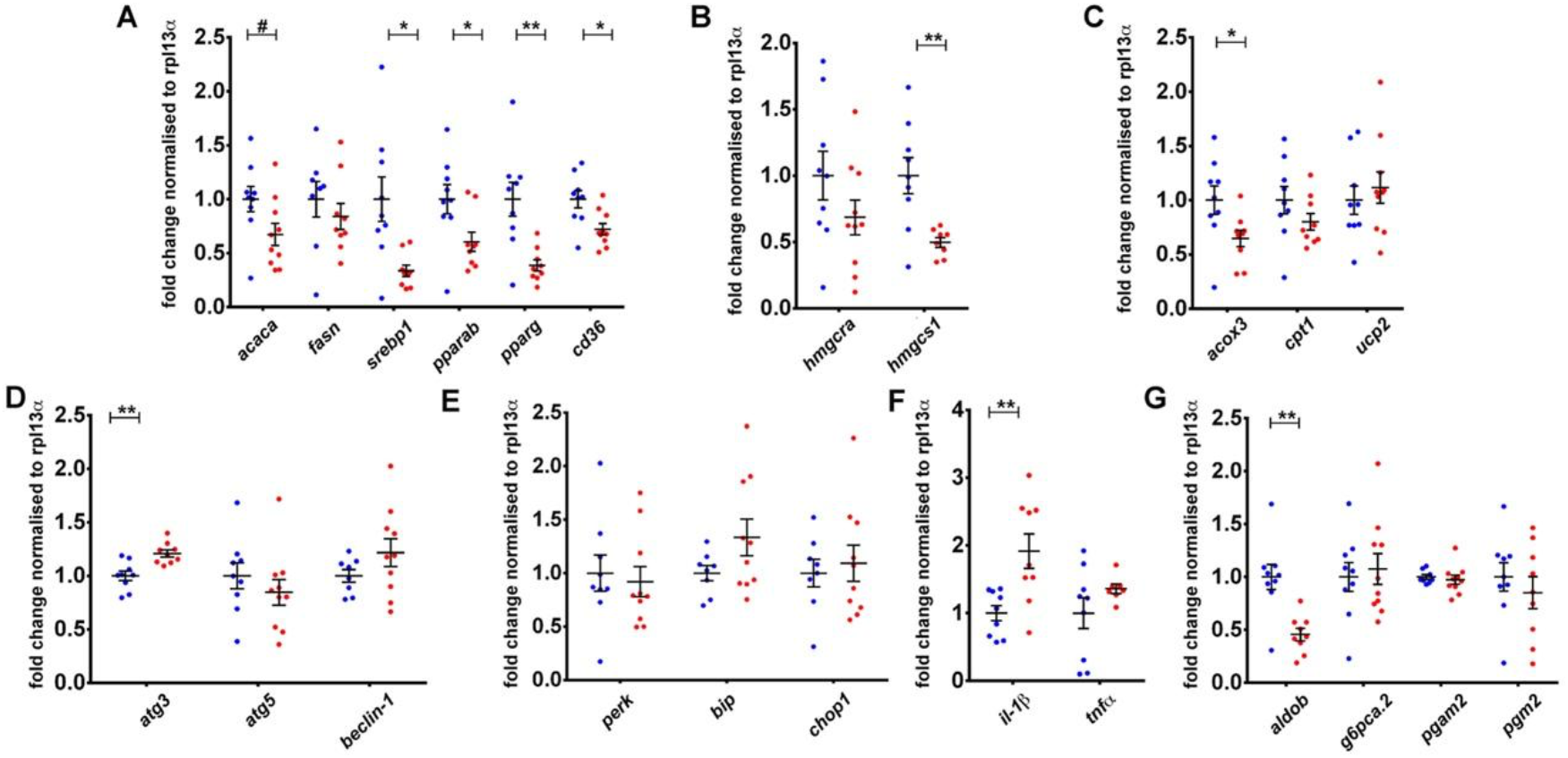
Gene expression changes in IL6-OE larvae. qRT-PCR of genes involved in lipogenesis, cholesterol metabolism, autophagy, unfolded protein response, inflammation and glycolysis. Fold change is normalized to *rpl13a.* Unpaired student’s t-test was applied to determine statistical significance. #p=0.05 *p<0.05, **p<0.005.

In our previous study the adult liver of IL6-OE zebrafish had also shown down regulation of autophagy genes. In contrast, we observed a slight but significant upregulation of *atg3* and no significant change in *atg5* and *beclin1* in the IL6-OE larvae (Fig 2D). Similarly, unfolded protein response (UPR) genes *perk* and *dnajc3* were reported to be down regulated in the adult IL6-OE liver(9), while we did not find any difference in expression of these genes at larval stage (Fig. 2E). We observed a significant increase in the inflammatory marker *IL1-beta* in larvae but not in adult liver, however no change was observed in *tnf-alpha* expression in either (Fig 2F). These differences may be attributed to the fact that unlike the adult, the larval gene expression was of the whole larvae and not specifically of the liver. Our adult transcriptomics and metabolomics study had revealed a dysregulation of glycolysis and gluconeogenesis genes in the IL6-OE liver(9). In the IL6-OE larvae we found that *aldob* expression was significantly downregulated reminiscent of the IL6-OE adult liver (Fig 2G). *ALDOB* deficiency is also known to lead to hepatic fat accumulation in patients (7).

### IL6 overexpression leads to increase in saturated triacylglycerols and a decrease in unsaturated triacylglycerols in adult male liver

Since we observed hepatic fat accumulation in the larval IL6-OE we compared the lipid composition of control and IL6-OE zebrafish larvae. We performed a targeted multiple reaction monitoring (MRM) lipidomics analysis of 7dpf larvae of control (n=5, each sample had 20 larvae) and IL6-OE (n=5). Lipids were extracted as described in the methods section and were detected and quantified using LC-MS/MS platform. We performed principal component analysis (PCA) using the online Metaboanalyst 4.0 tool (17). We observed that the control samples and the IL6-OE samples cluster separately from each other and within their own group (Fig. 3A). We identified 445 lipid species at a cut-off of 30% CV and count >6. Using a minimum fold change cutoff of 1.3 and of p<0.05 we identified 49 lipids out of the 445 detected that were differentially present in the IL6-OE larvae compared to control (Fig. 3B). Around 40 lipid species, predominantly phospholipids and some triacylglycerols (TAG), were more in IL6-OE larvae compared to control. We also identified nine lipids that were down regulated in the IL6-OE larvae compared to the control (Fig. 3B and Supplementary Information Fig. S1).

**Figure 3.**
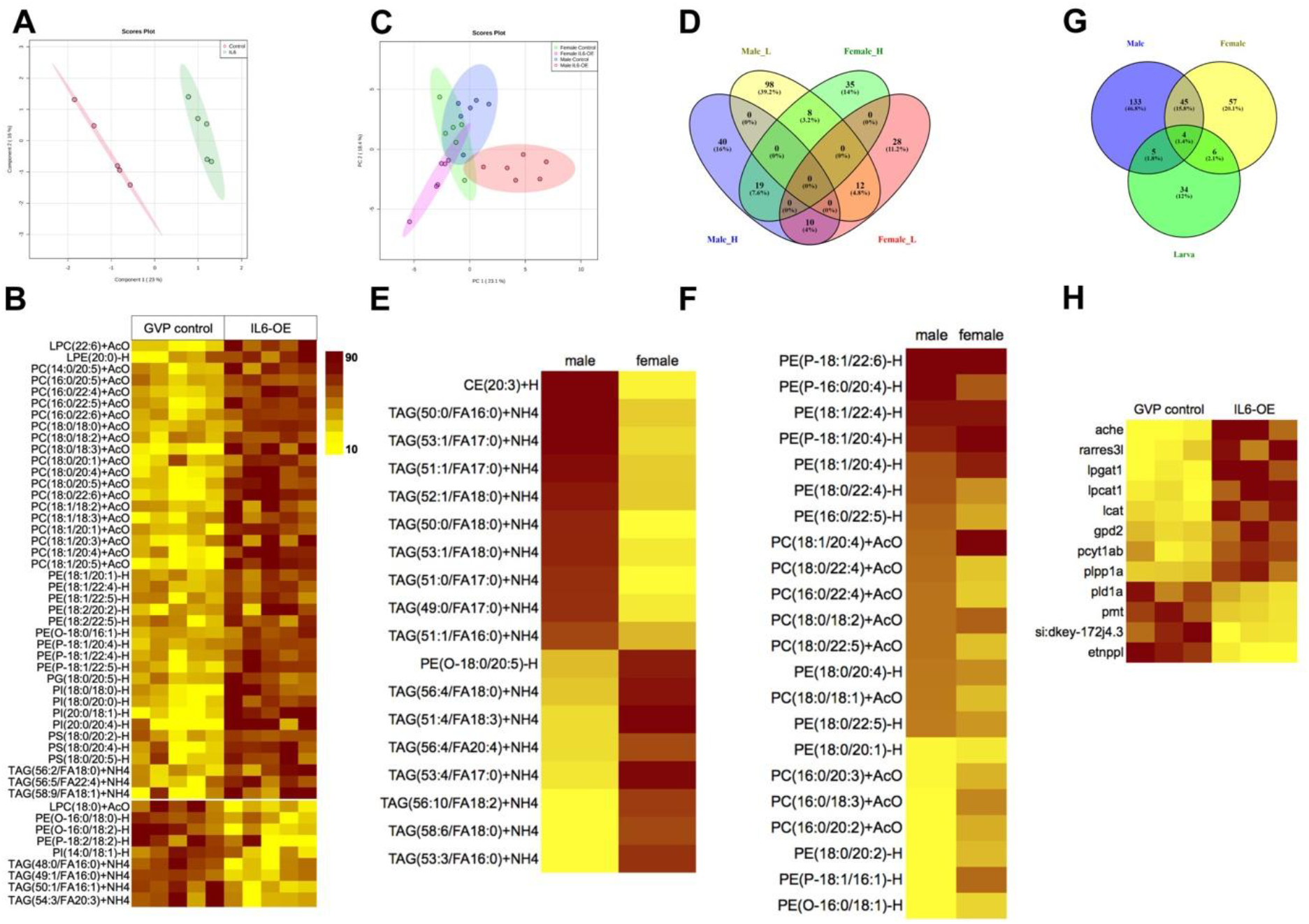
Lipidomics study reveals IL6-OE accumulates more saturated lipids. (A) PCA plot showing differences in lipid profile of control and IL6-OE larvae. (B) Lipids differentially present in IL6 overexpressed larvae. (C) PCA plot analysis of adult IL6-OE and control liver. (D) Venn diagram showing changes in lipid species between different experimental groups. (E) Selected lipids showing opposite patterns of accumulation in adult male and female IL6-OE livers. (F) Phospholipids differentially present in adult male and female IL6-OE liver. G) Venn diagram showing changes in lipid species common to IL6-OE larvae and adult. H) Expression of genes involved in lipid metabolism from RNA sequencing analysis. (B, E, F) Fold change of each sample was normalized against the average of GVP control and IL6-OE respectively. Heatmap was created according to the percentile value.

In the adult zebrafish, we had previously reported hepatic lipid accumulation selectively in the male and not female IL6-OE (9). To identify the differentially accumulating lipid species, we performed lipidomics profiling on 24 adult liver samples from 4 different groups: male control (n=6), male IL6-OE (n=6), female control (n=6), and female IL6-OE (n=6). PCA analysis of all the samples showed that they clustered distinctly into four separate groups (Fig 3C). In the overall comparison, 469 lipids were identified to be present at cut-off of 30% CV and count >6. Heat map analysis of these 469 showed dramatic changes in the lipid profile in the IL6-OE adults compared to sex specific controls (Supplementary Information Fig. S2). With a minimum cut-off of 1.3-fold change and p<0.05 we found 187 lipids differentially present (69 elevated and 118 reduced) in IL6-OE males and 112 lipids differentially present (62 elevated and 50 reduced) in the IL6-OE females (Fig. 3D). Among these were 18 lipid species that exhibited contrasting abundance in the male and female IL6-OE livers (Fig. 3E). Of these 18 lipids, 16 were TAGs; 9 TAGs that were up regulated in the male IL6-OE liver contained saturated and monounsaturated fatty acids (FA) and the 7 TAGs that were up regulated in the female IL6-OE contained poly-unsaturated FA (Fig. 3E). We also found a number of phospholipid species that were differentially expressed specifically in the male IL6-OE liver (Fig. 3F). Among the many differentially present lipids in males, we found 3 phosphatidylcholine species [PC(18:1/20:4)], PC(16:0/22:4)], and PC(18:0/18:2)] and 2 phosphatidylethanolamine species [PE(18:1/22:4)], and PE(P-18:1/20:4)] that were up regulated both in the IL6-OE larva and the adult male IL6-OE liver (Fig. 3G and Supplementary Information Fig. S3A). These results indicate that IL6 overexpression causes accumulation of phospholipids and more saturated triglycerides.

Since, there are significant changes in the lipid content of the control and adult IL6-OE zebrafish liver, we looked for lipid metabolism genes in our adult liver transcriptomics data. Glycerophospholipid metabolism was one of the important pathways that was differentially regulated in the IL6-OE zebrafish (Supplementary Information Fig. S3B) (9). We also found significant changes in the expression of a number of genes in lipid pathway (Fig. 3H) indicating large changes in the metabolic program of the IL6-OE liver (Supplementary Information Fig. S3B).

### Rosiglitazone ameliorates hepatic fat accumulation in IL6-OE larvae

We performed a small targeted screen of small molecules to identify compounds that could reverse or ameliorate the hepatic lipid accumulation in the IL6-OE zebrafish larvae. Since, the IL6-induced lipid accumulation is likely mediated by JAK-STAT3 signaling pathway induced by the IL6 receptor, we selected two known inhibitors of STAT3 signaling; LLL12 and LMT28(18,19). From our adult and larval studies we have observed a consistent down regulation of PPAR-alpha and PPAR-gamma expression in the IL6-OE zebrafish(9) (Fig. 2A). Thus, we selected three compounds: Wy14643, a PPAR-alpha agonist and Rosiglitazone and Pioglitazone, agonists of PPAR-gamma for the screen(20–22).

Briefly, 5dpf larvae were treated with 10μM concentration of the small molecules for 48 hours. Animals were fixed in paraformaldehyde at 7dpf and were stained with ORO to detect lipid accumulation (Fig 4A). When compared to vehicle-treated IL6-OE larvae, Rosiglitazone and Pioglitazone treatment appeared to reduce the lipid accumulation in the zebrafish liver. The other three compounds did not show any obvious difference in the ORO staining. Upon repeating the treatment, Rosiglitazone (RGT) consistently reduced the ORO staining in the liver. We then tested increasing concentrations of RGT from 5μM to 25μM on the 5dpf zebrafish larvae and found that there was a more pronounced reduction of ORO staining at 25μM compared to all other concentrations (Fig. 4B). 5dpf and 8dpf larvae were treated with 25μM RGT for 48 hours. The 10dpf larvae showed a more striking reduction in ORO staining compared to 7dpf (Fig. 4C). The treatment however had no effect on the liver size. Adult male IL6-OE were treated with 25μM RGT in water daily for 5days. Liver was fixed and sectioned for ORO staining. Although compared to DMSO-treated IL6-OE liver sections, treatment with RGT did appear to have an effect on lipid staining, it was not consistent (Fig. 4C). We also quantified the total triacylglycerol in adult liver by thin layer chromatography. However, the difference between control and IL6-OE was not significant enough to derive any conclusion (Supplementary information Fig. S4).

**Figure 4.**
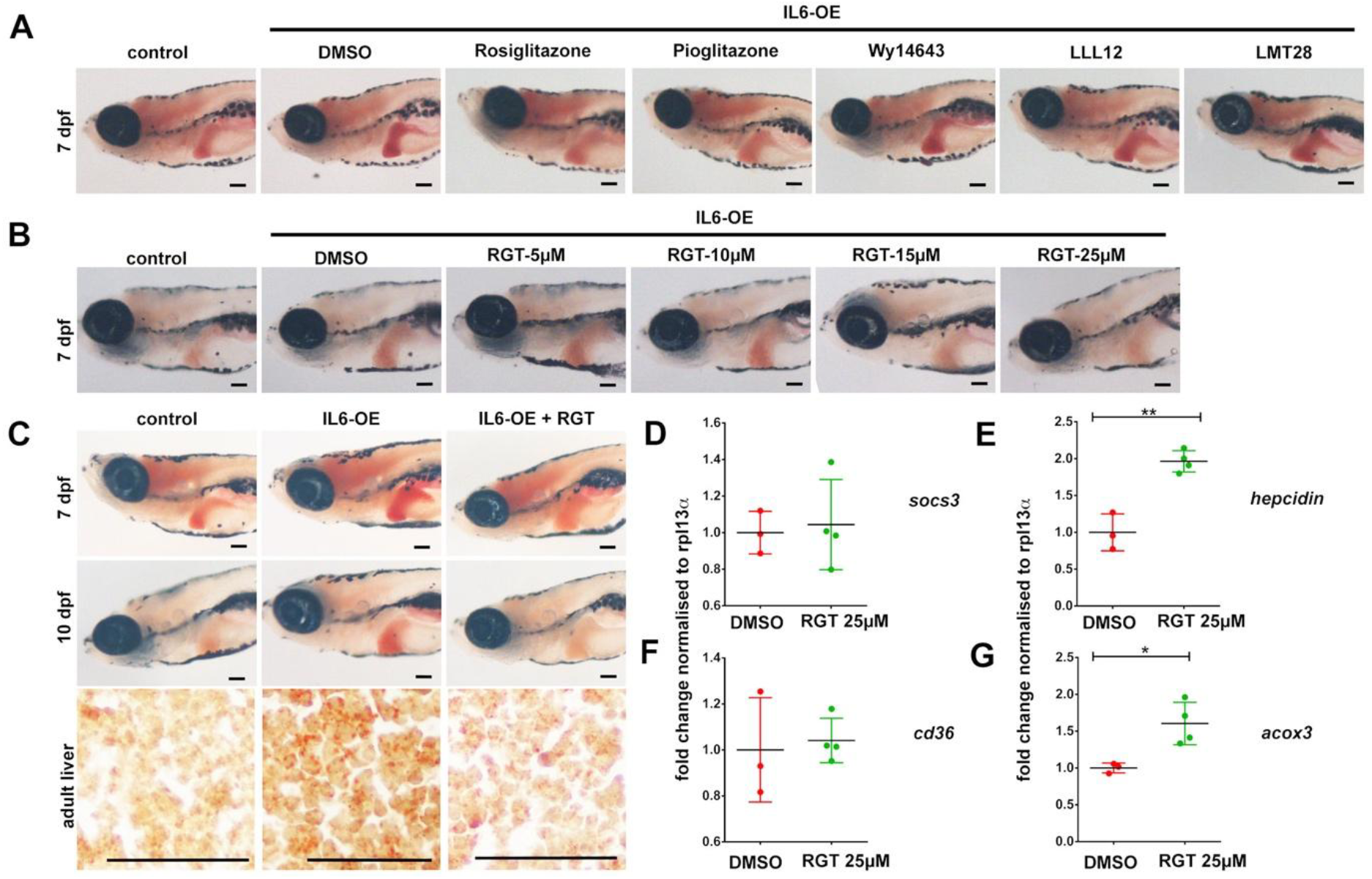
Rosiglitazone treatment ameliorates fat accumulation in IL6-OE zebrafish. A) Oil red O staining of 7 dpf larvae shows that treatment with 10μM Rosiglitazone and Pioglitazone reduces fat accumulation in IL6-OE. B) Oil red O staining of 7 dpf larvae shows that treatment with 25μM Rosiglitazone showed a pronounced reduction in fat accumulation in IL6-OE. C) 25μM Rosiglitazone treatment had more striking reduction in fat accumulation at 10dpf in comparison to 7dpf. Oil red O staining of adult liver sections shows a marginal but unremarkable reduction of fat accumulation in Rosiglitazone treated IL6-OE liver. (D-G) real time PCR of 10 dpf larvae treated with Rosiglitazone shows no effect on expression of *socs3* (D) and *cd36* (F). However, there is significant induction of hepcidin (E) and *acox3* (G). Unpaired student’s t-test was applied to determine statistical significance. *p<0.05, **p<0.01. Scale bar is 10μm.

To investigate the mechanism by which RGT ameliorates hepatic fat accumulation in the larvae we examined expression of genes involved in IL6 signaling, lipid metabolism and peroxisomal β-oxidation. We did not find any difference in expression of *socs3* indicating RGT does not modulate IL6 signaling (Fig. 4D). Surprisingly *hepcidin* expression was upregulated two-fold; the significance of this is not clear to us (Fig. 4F). *cd36* is a fatty acid uptake receptor and we found no change in its expression upon RGT treatment (Fig. 4E). However, there was a significant upregulation of acox3, the acetyl coenzyme A oxidase, that causes desaturation of fatty acids in the peroxisomes(23). *acox3* expression was significantly reduced in the IL6-OE fish compared to controls (Fig. 2C) and this effect appears to be reversed by RGT (Fig. 4G). Thus, these observations suggest that activation of PPAR-gamma signaling enhances peroxisomal β-oxidation through *acox3* upregulation perhaps affecting the saturation of fatty acids in the liver.

### Rosiglitazone increases unsaturated TAGs and reduces saturated TAGs

To test the status of various lipids in the RGT treated IL6-OE fish, we performed lipidomics on the larvae. We found 61 lipids that were differentially present in IL6-OE larvae treated with RGT compared to DMSO. Of the 35 differentially regulated TAGs observed in the larvae, which are predominantly polyunsaturated, 15 were downregulated and 20 were upregulated upon treatment (Fig. 5A). We also observed the changes in phospholipid composition upon RGT treatment in larvae.

**Figure 5.**
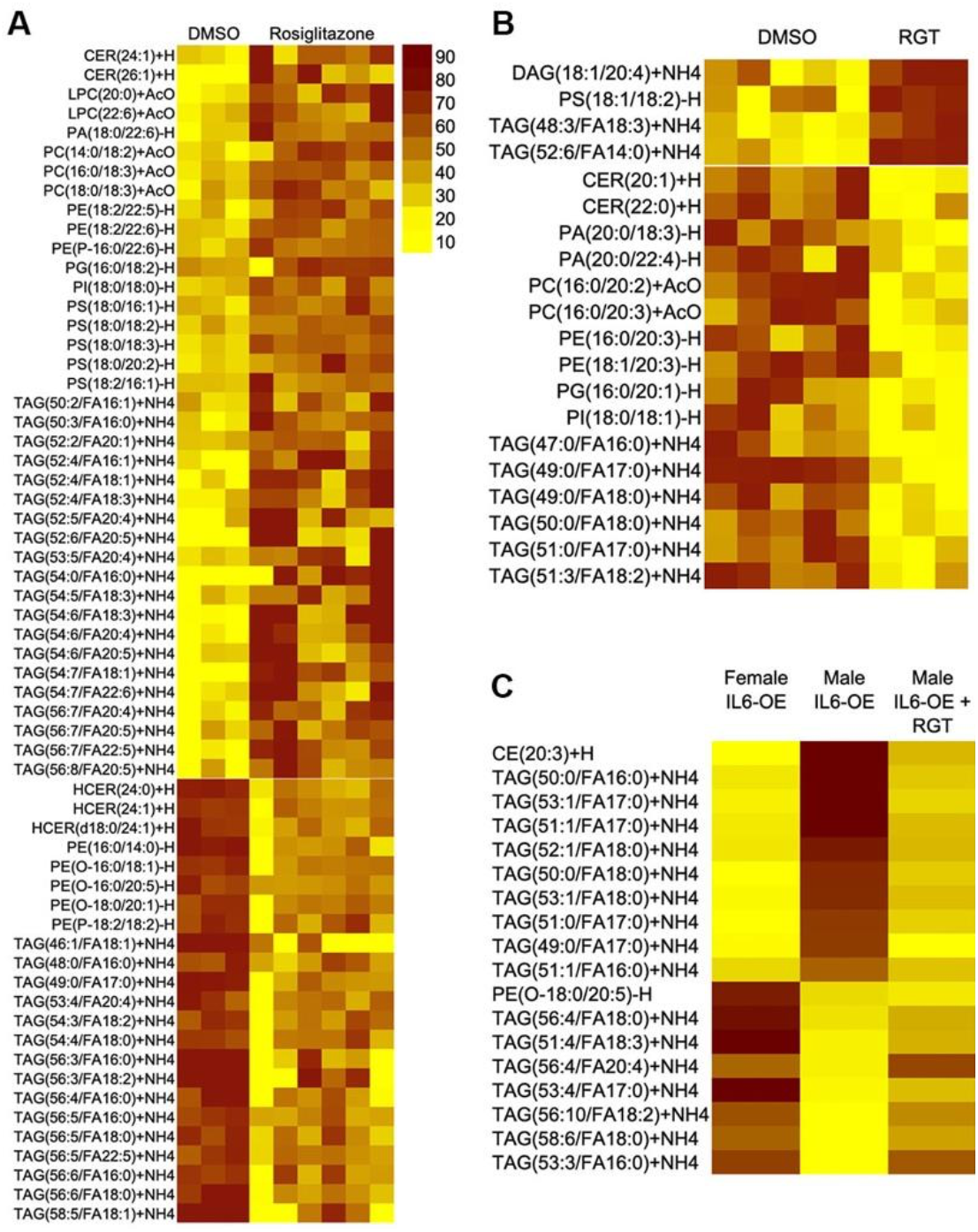
Rosiglitazone treatment increases unsaturated TAGs and reduces saturated TAGs. A) 61 lipids were differentially present in the whole larvae after RGT treatment. Of these 38 lipids showed upregulation and 23 lipids showed downregulation upon RGT treatment. B) Rosiglitazone treatment of adult IL6-OE showed down regulation of hepatic saturated TAGs C) 18 lipids with opposite patterns in male and female IL6-OE are reversed upon RGT treatment. Fold change of each sample was normalized against the average of GVP control and IL6-OE respectively. Heatmap was created according to the percentile value.

Adult IL6-OE were exposed to 25μM RGT in water daily for 5 days and lipidomics analysis of liver was performed. The TAGs downregulated by RGT treatment in the adult liver predominantly contained saturated fatty acids (Fig. 5B). In addition, a number of phospholipids also showed differential levels in the DMSO and RGT treated adult liver (Fig. 5B). Previously we had identified 18 lipid species that were regulated in an opposite manner in the adult IL6-OE male and female livers (Fig. 3E). These contained saturated TAGs that accumulated in the male IL6-OE compared to female IL6-OE liver and unsaturated TAGs that accumulated more in female than male IL6-OE. We found that RGT treatment of male IL6-OE livers led to a reversal in trend of most of these 18 lipids such that the RGT-treated male liver lipid-profile appeared to resemble the female lipid pattern (Fig. 5C).

## Discussion

Fatty liver is most commonly associated with obesity and most animal models use high fat or high fructose diet to create fatty liver condition. Our previous studies demonstrated that chronic systemic exposure to IL6 leads to hepatic fat accumulation in adult zebrafish (9). This phenomenon however was specific to male zebrafish as the female liver did not accumulate lipids in spite of having similar exposure to IL6. However, it was not clear if the hepatic accumulation in the IL6-OE was equivalent to the fatty liver in NAFLD in humans. In the present study we performed a lipidomics analysis on the IL6-OE zebrafish to determine the types of lipids present in the liver of male and female liver.

We found that in adult IL6-OE fish, the male liver accumulated more saturated lipids especially TAGs and that this was accompanied by a down regulation of unsaturated lipids. The female lipidomic profile reflected an opposite trend with increase in unsaturated lipids and a down regulation of saturated lipids in the IL6-OE compared to controls. This suggests a protective mechanism of action in the females, which needs further investigation. Lipidomic analysis of whole larvae found a number of lipids to be perturbed in the IL6-OE compared to controls. Here also we found accumulation of more saturated lipids than unsaturated. Studies on NAFLD patient livers have shown a shift from unsaturated to saturated lipid profile (24,25). Thus, our zebrafish chronic inflammation model appears to resemble human NAFLD pathology.

Previous gene expression studies of the adult IL6-OE fish had revealed major changes in lipid and glucose metabolism(9). In the present study we discovered that these changes were also evident in the IL6-OE larvae reinforcing that chronic systemic IL6 exposure causes re-wiring of the metabolic pathways in liver towards lipid synthesis. In our adult zebrafish liver transcriptomics studies, we had observed a down regulation of PPAR signaling in the IL6-OE (9). This was reinforced in the IL6-OE larvae as well. This indicated that IL6-mediated accumulation of saturated fatty acids in the liver might be mediated through downregulation of PPAR signaling. Lean NAFLD patients also show a similar lipid profile with a decrease in unsaturated phospholipids and triglycerides compared to controls (11). Our targeted chemical genetics approach led to the identification of PPAR-gamma agonist Rosiglitazone (RGT), a compound that could reduce the hepatic fat accumulation in the larvae. Lipid profiling of livers from RGT treated fish showed a reduction in saturated fatty acids and an increase in unsaturated fatty acids in the male IL6-OE fish. We also found that RGT treatment led to the upregulation of *acox3* gene expression, an enzyme involved in peroxisomal beta-oxidation and unsaturation of fatty acids. Pioglitazone which is a PPAR-γ activator, has shown reversal of NASH and fibrosis in phase 4 clinical trials (26,27) suggesting similar disease mechanisms to be operative in the NAFLD patients and our zebrafish IL6-OE model.

Our study reveals that chronic systemic exposure to IL6 leads to a change in fatty acid composition of the liver in adult and larval zebrafish. Further, it suggests that activation of the PPAR-gamma pathway can increase acox3 expression and lead to desaturation of fatty acids. Thus PPAR-gamma could serve as a useful target while developing therapeutics for NAFLD. More importantly, we have demonstrated that our diet and obesity independent model of NAFLD in zebrafish is responsive to similar drugs as has been found in pre-clinical studies on NAFLD models and on NAFLD patients. Thus, the IL6-OE model can serve as an effective model to study obesity-independent NAFLD as well as an excellent and tractable system for chemical screening and drug discovery.

## Materials and Methods

### Zebrafish line and maintenance

Zebrafish *(Danio rerio)* were bred and maintained under standard conditions and temperature at 28.5°C. Larvae were grown and fed with paramecia for one month followed by regular feed thereafter(28). Handling of larvae and adult zebrafish was in strict accordance with good animal practices as described by the Committee for the Purpose of Control and Supervision of Experiments on Animals (CPCSEA), Government of India. All the experiments were approved by The Institutional Animal Ethics Committee (IAEC) of the CSIR-Institute of Genomics and Integrative Biology, New Delhi, India.

Zebrafish transgenic lines that were used in this study - *Tg (pCH-cmlc2:GVP)* and *Tg(pCH-cmlc2:GVP::pBH-UAS:IL6)(29).*

### Whole mount RNA in situ hybridization

RNA in situ hybridization was performed as described (30). Digoxigenin-labeled antisense riboprobe against human IL6 gene was used as described previously(9).

### Quantitative Real Time PCR

Real time PCR and statistical analysis was performed on larval and liver tissue as described (9). The primers used in this study has been previously described (9).

### Histology and staining

Zebrafish liver tissues were fixed in 4% paraformaldehyde. Cryo-sections of 7μm thickness were used for Oil Red O staining. For oil red o staining larvae were fixed in 4 % PFA at 4°C overnight. Washed thrice with PBS and incubated in preheated 90% propylene glycol at 50°C for 1hr. Further, Propylene glycol was replaced with ORO and incubated at 50°C for 1hr followed by 3 washes in 70% PG for 30 min each. Then finally washed with PBS and larvae were imaged. Nikon upright microscope (model E200) was used for Oil Red O and H&E sections.

### Lipid extraction and thin layer chromatography

Total lipids were extracted from liver tissue and whole larva using the Modified Bligh and Dyer Method(31). The dried lipids were further re-suspended in chloroform: methanol (2:1) and were loaded on a thin-layer chromatography plate. Lipids are allowed to separate on TLC plates in the solvent system hexane: diethyl ether: acetic acid (70:30:1). TLC plates were sprayed using 10% copper sulphate (w/v) in 8% phosphoric acid (v/v) solution. Further, the plates are dried at 150°C for signal development. Quantification of un-labeled lipid spots was done using ImageJ® software(32).

### Drug treatment larvae and adult

Zebrafish IL6-OE adults were bred and embryos were collected. Embryos were allowed to grow till 5dpf at 28°C. Larvae were then treated with small molecules or DMSO from 5dpf to 7dpf at 28°C.

For chemical treatment of adults, small molecules were dissolved in the appropriate amount of water to make a final concentration of 25μM. Zebrafish were kept in DMSO or small molecules dissolved in water for 5 days. These adults were then sacrificed to collect liver tissue on the 6th day for further processing.

### Lipidomics

The lipids were extracted by a modified Bligh and Dyer method(31), using a mixture of dichloromethane/methanol/water (2:2:1 v/v). The lyophilized extracts were reconstituted in 100 % ethanol and subjected to LC-MS run. Lipidomics was done using An ExionLC™ Exion LC system with a Waters AQUITY UPLC BEH HILIC XBridge Amide column (3.5 μm, 4.6 x 150 mm). Binary gradient– Buffer A (95% acetonitrile in 10mM ammonium acetate, pH-8.2) and Buffer B (50% acetonitrile in 10mM ammonium acetate, pH-8.2) were used for chromatographic separation with a flow rate was adjusted toof 0.7 ml/minute. A theoretical MRM was generated using LIPIDMAPS for identification and relative quantification of lipids. Sciex QTRAP 6500+ system in low mass range with Electrospray Ionization (ESI) probe was used for identification and relative quantification of lipid species(33).

### RNA sequencing and Statistical analysis

RNA sequencing analysis of previously published data from our lab(7) (Gene expression project Accession: PRJNA638724) was done using standard pipeline. Differential gene expression was analyzed using fold change cutoff of 2. The genes that showed significance of <0.05 were used further for analysis.

## Contribution

Manoj K Singh: Study concept and design, acquisition of data, analysis and interpretation of data and drafting of manuscript.

Rohit Yadav: Study design, acquisition of data, analysis and interpretation of data and drafting of manuscript.

Akash Kumar Bhaskar: acquisition and analysis of lipidomics data.

Shantanu Sengupta: interpretation of lipidomics data.

Chetana Sachidanandan: Study concept and design, analysis and interpretation of data, drafting of manuscript, obtained funding.

## Funding information

This work was supported by the Council of Scientific and Industrial Research (CSIR), New Delhi [BSC0124, MLP1801]. Manoj Kumar Singh and Rohit Yadav was supported by Council of Scientific and Industrial Research (CSIR) research fellowship. The funders had no role in study design, data collection and analysis, decision to publish, or preparation of the article.

## Declaration of competing interest

The authors declare that they have no conflict of interest.

## Acknowledgments

We appreciate the help of Narendra Kumar for zebrafish lab maintenance. We acknowledge Dilip Menon for discussions on lipidomics experiments. We also acknowledge Ahmed Mobeen, Anasuya Bhargava and Surbhi for discussions and help during the study.

## Notes

### Competing Interest Statement

The authors have declared no competing interest.

