## Supplementary figure for "A diet-independent zebrafish model for NAFLD recapitulates patient lipid profiles and offers a system for small molecule screening"

### Supplementary Information

**Rosiglitazone, a PPAR-gamma agonist, modulates hepatic fat accumulation in an inflammation-induced fatty liver model in zebrafish**

Manoj K Singh[1,2,*], Rohit Yadav[1,2], Akash Bhaskar[1,2], Shantanu Sengupta[1,2], Chetana Sachidanandan[1,2,*]

S 1

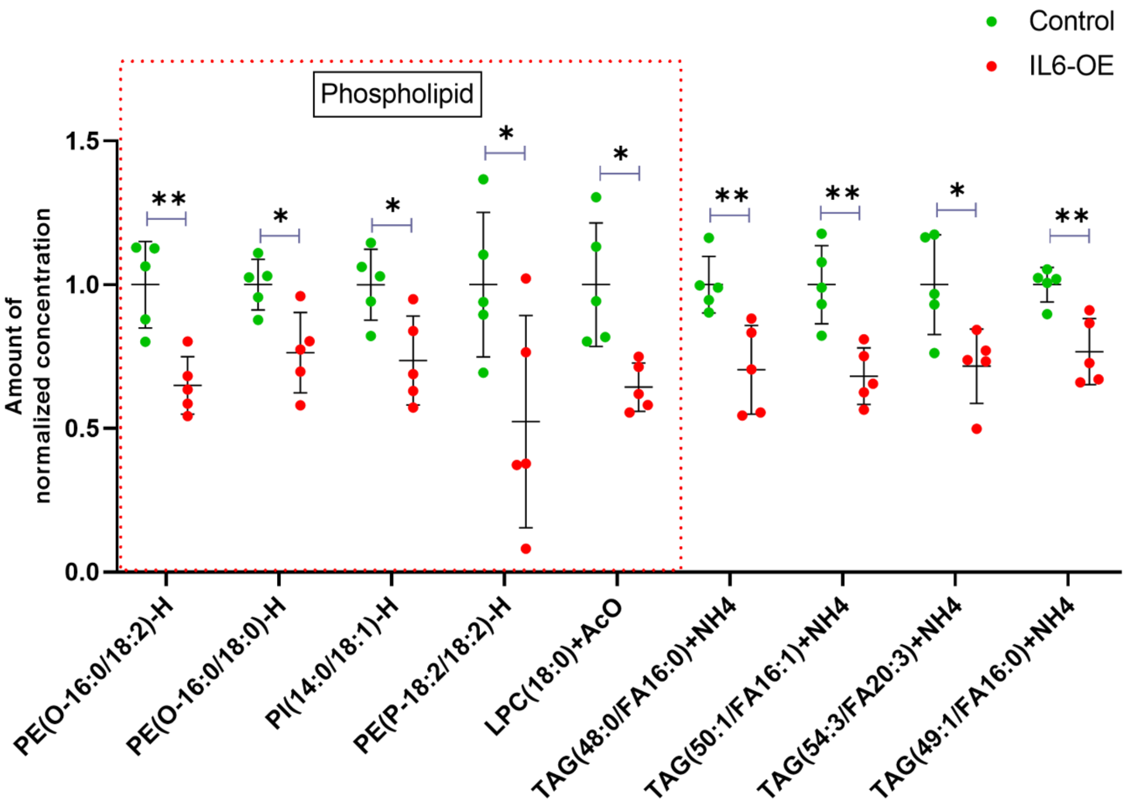


**Supplementary Figure 1. Phospholipids downregulated in IL6-OE larvae.** Lipidomics study of IL6-OE larvae showed 49 lipids to be differentially regulated in comparison to control. Out of 49 lipids we found 9 lipids were downregulated which majorly constitute phospholipids. *p<0.05, **p<0.01. Dotted red square shows downregulated phospholipid.

S 2

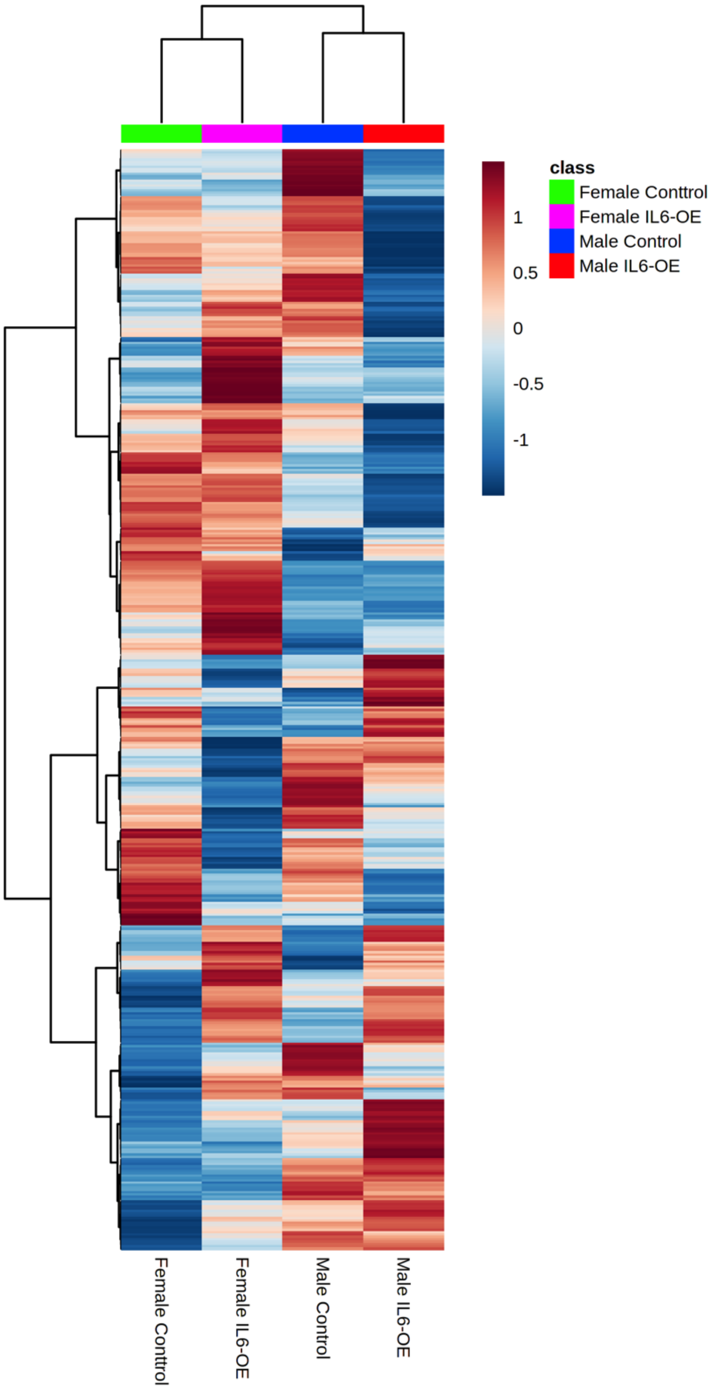


**Supplementary Figure 2. Differentially present lipids in IL6-OE adults.** Lipidomics analysis of adult IL6-OE reveals 469 differentially present lipids at a cut-off of 30% CV and count of 6. Heatmap of these 469 showed dramatic changes in the lipid profile in the IL6-OE compared to sex specific controls.

S 3

A

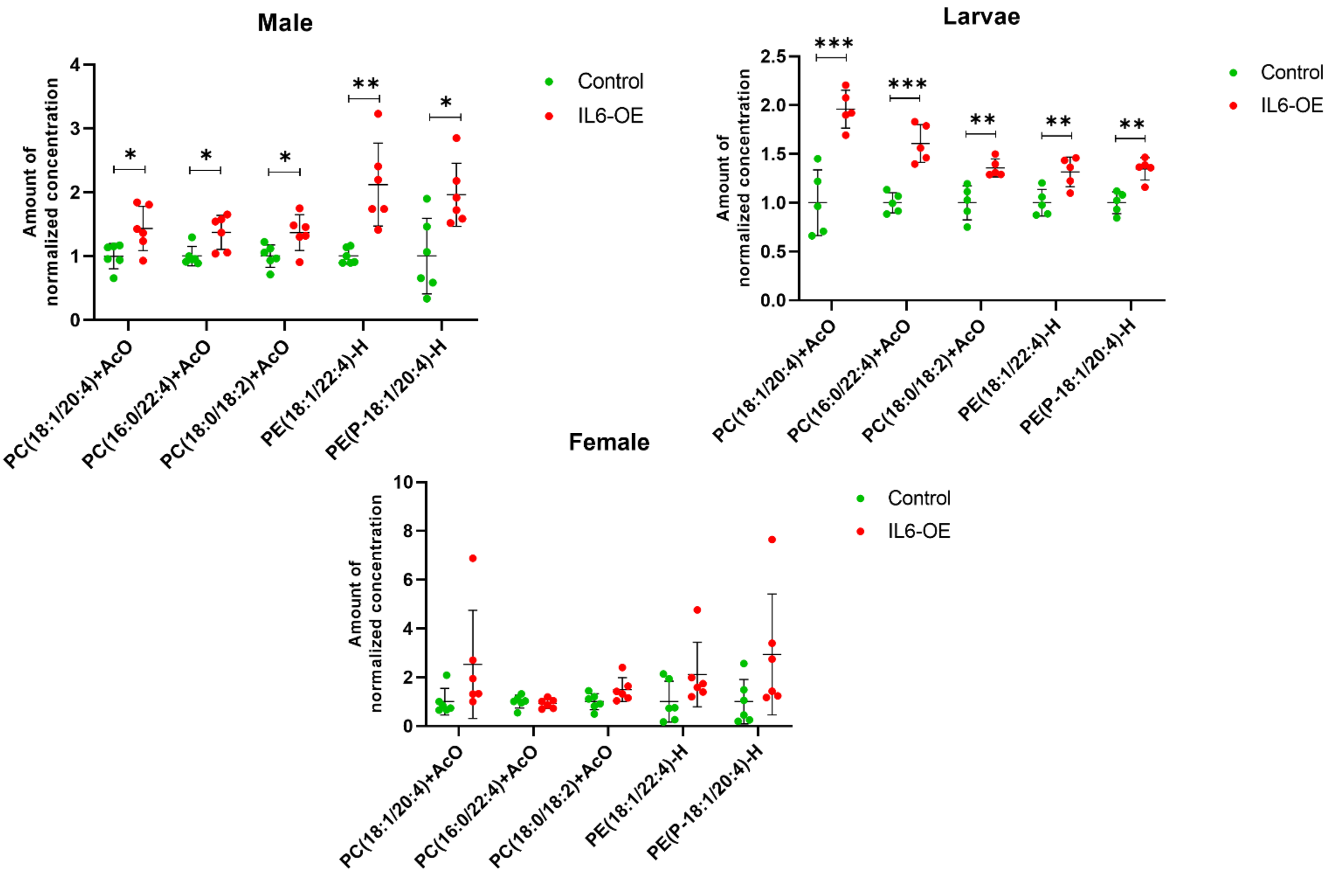
Male
Larvae
Female
Amount of normalized concentration
Control
IL6-OE
PC(18:1/20:4)+AcO
PC(16:0/22:4)+AcO
PC(18:0/18:2)+AcO
PE(18:1/22:4)-H
PE(P-18:1/20:4)-H

B

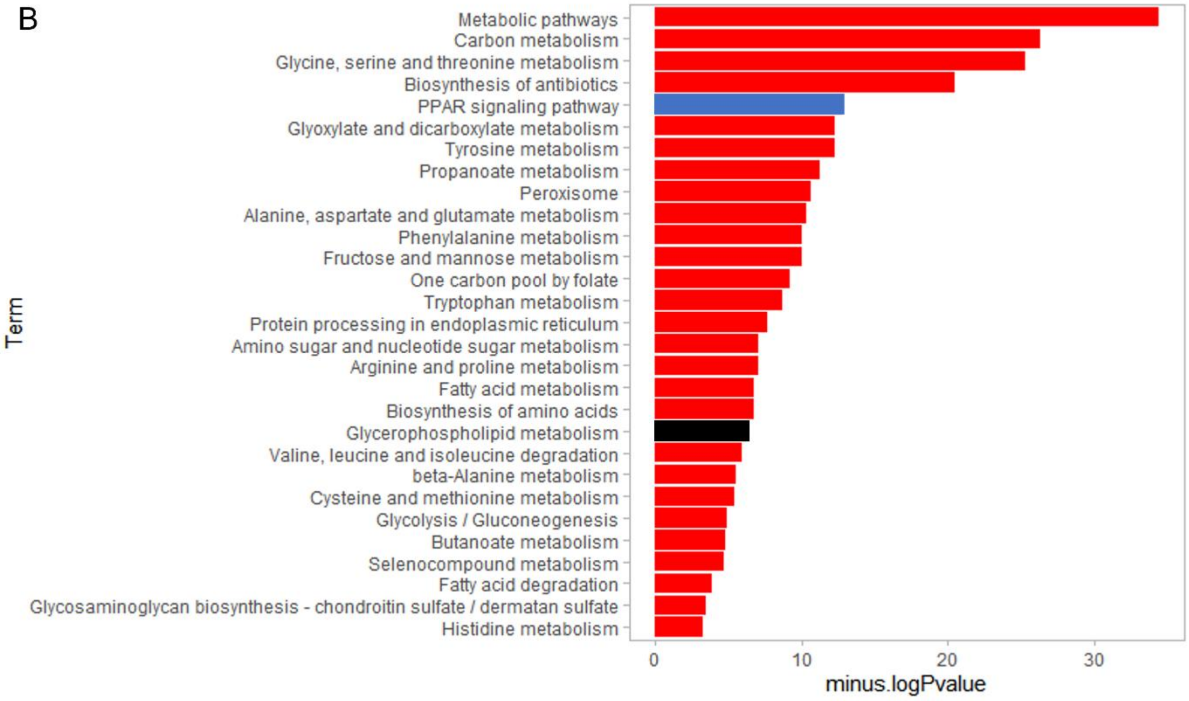
Term
Metabolic pathways
Carbon metabolism
Glycine, serine and threonine metabolism
Biosynthesis of antibiotics
PPAR signaling pathway
Glyoxylate and dicarboxylate metabolism
Tyrosine metabolism
Propanoate metabolism
Peroxisome
Alanine, aspartate and glutamate metabolism
Phenylalanine metabolism
Fructose and mannose metabolism
One carbon pool by folate
Tryptophan metabolism
Protein processing in endoplasmic reticulum
Amino sugar and nucleotide sugar metabolism
Arginine and proline metabolism
Fatty acid metabolism
Biosynthesis of amino acids
Glycerophospholipid metabolism
Valine, leucine and isoleucine degradation
beta-Alanine metabolism
Cysteine and methionine metabolism
Glycolysis / Gluconeogenesis
Butanoate metabolism
Selenocompound metabolism
Fatty acid degradation
Glycosaminoglycan biosynthesis - chondroitin sulfate / dermatan sulfate
Histidine metabolism
minus.logPvalue

**Supplementary Figure 3. Common phospholipids upregulated in adults and larvae.** A) Comparison of phospholipid profile of IL6-OE adult and larvae shows that 5 phospholipids to be commonly upregulated in adult male and larvae while no significant change was observed in female IL6-OE adult. Among these 3 were phosphatidylcholine species and 2 were phosphatidylethanolamine species of lipids. *$p<0.05$, **$p<0.01$, ***$p<0.001$. B) IL6-OE adult male showed glycerophospholipid metabolism genes to be differentially regulated. We also found significant changes in genes involved in lipid metabolism pathways like- PPAR signaling. Blue box in the figure represents PPAR signaling genes and black box represent glycerophospholipid metabolism genes.

S 4

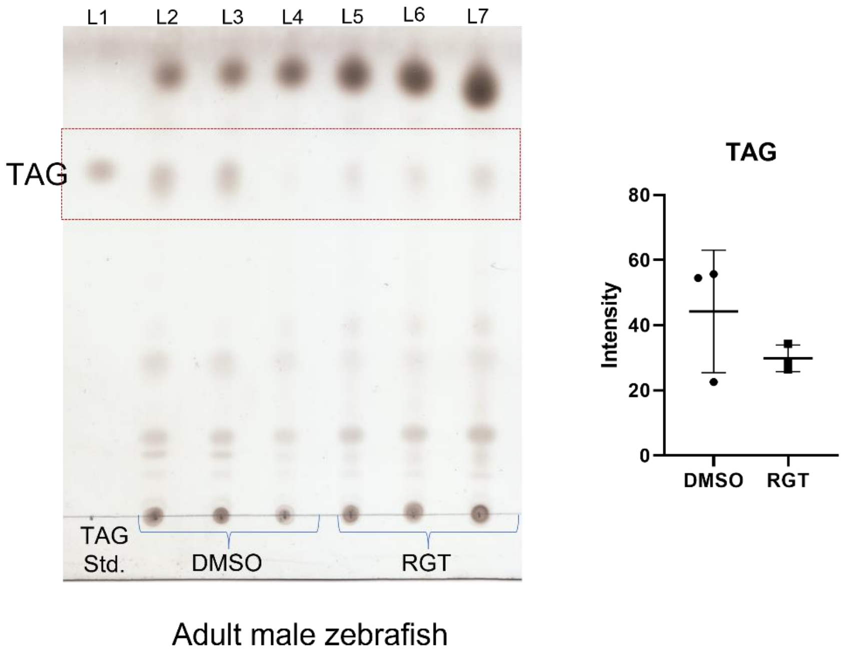


**Supplementary Figure 4. Quantification of total hepatic triacylglycerol using TLC.** A) L1- triacylglycerol standard, L2-L4 - DMSO treated adult male IL6-OE liver, Lane L5-L7 - Rosiglitazone treated adult male IL6-OE liver. B) Quantified TAG using ImageJ software. Total triglyceride level shows a trend of reduction in adult IL6-OE male after RGT treatment.
